# Genotype-by-Environment Interactions Affecting Heterosis in Maize

**DOI:** 10.1101/131342

**Authors:** Zhi Li, Lisa Coffey, Jacob Garfin, Nathan D. Miller, Michael R. White, Edgar P. Spalding, Natalia de Leon, Shawn M. Kaeppler, Patrick S. Schnable, Nathan M. Springer, Candice N. Hirsch

## Abstract

The environment can influence heterosis, the phenomena in which the offspring of two inbred parents exhibits phenotypic performance beyond the inbred parents for specific traits. In this study we measured 25 traits in a set of 47 maize hybrids and their inbred parents grown in 16 different environments, and each had varying levels of average productivity. By quantifying 25 vegetative and reproductive traits across the life cycle we were able to analyze interactions between the environment and multiple distinct instances of heterosis. The magnitude and rank among hybrids of better-parent heterosis (BPH) varied for the different traits and environments. Across the traits, a higher within plot variance was observed for inbred lines compared to hybrids. However, for most traits, variance across environments was not significantly different for inbred lines compared to hybrids. Further, for many traits the correlations of BPH to hybrid performance and BPH to better parent performance were of comparable magnitude. These results indicate that inbreds and hybrids are showing similar trends in environmental response and are both contribute to genotype-by-environment interactions for heterosis. This study highlights that degree of heterosis is not an inherent trait of a specific hybrid, but varies depending on the trait measured and the environment where that trait is measured. Studies that attempt to correlate molecular processes with heterosis are hindered by the fact that heterosis is not a consistent attribute of a specific hybrid.

## INTRODUCTION

Heterosis, or hybrid vigor, refers to the phenomena in which the offspring of two inbred parents exhibits phenotypic performance beyond the mid-parent or best parent used to generate the hybrid. Heterosis has been observed in many plant and animal species (Janick 1998; Melchinger and Gumber 1998). Notably, the heterosis of mules (the ability to perform more work with fewer resources) was widely utilized in agriculture prior to mechanization (Troyer 2006). Inbreeding depression and heterosis in maize was initially documented by George H. Shull and Edward M. East (East 1908; Shull 1908; Shull 1909). The adoption of hybrid maize over open-pollinated varieties occurred remarkably fast due to improved yields, greater uniformity for machine harvesting, and increased durability under extreme abiotic stress. In just a four-year period of time, hybrid maize acreage went from less than 10% to over 90% in Iowa (Crow 1998). The widespread utilization of heterosis now shapes breeding programs for several agriculturally important species including maize and rice.

There is widespread interest in developing methods to characterize the molecular basis of heterosis, and to predict hybrid performance to increase the efficiency of hybrid breeding programs. Researchers have attempted to utilize genomic sequence (Riedelsheimer et al. 2012), RNA expression levels of genes (Melchinger and Gumber 1998; Frisch et al. 2010; Scholten and Thiemann 2013), sRNAs (Groszmann et al. 2011; Zhang et al. 2014), proteomic (Dahal et al. 2012), and metabolomic (Riedelsheimer et al. 2012; de Abreu et al. 2017) data to predict or dissect heterosis (Schnable and Springer 2013). While relationships have been identified using each of these data types, no data type is able to completely predict hybrid performance individually (Kaeppler 2012). Attempts to predict hybrid performance are complicated by the fact that heterosis levels vary for different traits within the same hybrid (Flint-Garcia et al. 2009).

Although plant breeders have noticed that hybrid genotypes are more stress tolerant than their inbred parents, there are few published reports to support this conclusion, particularly in environments with moderate rather than extreme levels of abiotic stress. In *Arabidopsis*, stress response gene expression networks have been shown to contribute to heterosis and the prediction of hybrid performance (Groszmann et al. 2015; Miller et al. 2015). While variation in levels of heterosis have been observed under different growing conditions, there are few studies that document changes in heterosis across diverse environmental conditions and traits.

In this study we measured 25 traits in 47 maize hybrids and their inbred parents that were grown in 16 different environments. The objective of the study was to document variation in estimates of heterosis across traits and environments. The results provide evidence that unravelling the molecular basis of heterosis is challenging because heterosis is not a fixed attribute of an individual across space, time, or environmental conditions.

## MATERIALS AND METHODS

### Germplasm

Eleven inbred lines were selected representing important founders in commercial maize breeding programs including DK3IIH6 (PI 564754), LH145 (PI 600959), LH185 (PI 576171), LH198 (PI 557563), LH82 (PI 601170), PHB47 (PI 601009), PHK56 (PI 543842), PHK76 (PI 601496), PHN46 (PI 543844), PHP02 (PI 601570), and a recent release W606S. These inbred lines represent multiple heterotic groups including Iodent (DK3IIH6, PHP02), Non-Stiff Stalk (LH185, LH82, PHK76, PHN46, PHK56, W606S), and Stiff Stalk (LH145, LH198, PHB47). These lines, with the exception of W606S, are all commercial inbred lines that have expired Plant Variety Protection certificates, and thus represent elite maize germplasm. These inbred lines were crossed in a partial diallel to generate 47 hybrid genotypes (see Table S1).

To evaluate genetic diversity between the parental lines used to generate the hybrid genotypes, genetic similarity between the parents was calculated using whole genome identity by state (Purcell et al. 2007) using 430,000 SNPs derived from RNA-sequencing (Hirsch et al. 2014).

### Field Evaluations

Trials containing single row plots (3.35 m long and 0.76 m apart) were planted in a total of 16 environments in Iowa, Minnesota, and Wisconsin in the summer of 2015. The 16 environments were defined by location (5 separate locations), and management practices within location (planting date; high (70,000 plants ha^-1^) and low (20,000 plants ha^-1^) plant density). Arlington, WI and Waseca, MN had high and low planting densities, representing a total of four environments. Curtiss, IA, Kelly, IA, and St. Paul, MN had a factorial of high and low planting density and early and late planting at each site, representing a total of 12 environments (see Table S2). Within each location/management environment there were two replications and hybrids were blocked separately from inbred lines within each replication.

Twelve vegetative traits were measured on six representative plants per plot. These traits included plant height at 14, 21, 28, 35, 42, 49, 56, and 63 days after planting (DAP) measured as the distance from the soil surface to the uppermost leaf tip when the leaves were pulled upright. Plant height at maturity was measured from the soil surface to the collar of the flag leaf, ear height at maturity was measured from the soil surface to the node subtending the uppermost ear. Leaf number above the ear, and leaf number (including senesced leaves) below the ear were counted after anthesis. Juvenile leaves were marked to allow leaf number including senesced leaves to be counted using previously described methods (Hirsch et al. 2014). Days to anthesis and days to silk were measured on a per-plot basis as the day on which approximately half of the plants in the plot were shedding pollen and the day on which half of the plants in the plot had exposed silks, respectively. Custom computer algorithms executed on Open Science Grid computational resources (Pordes et al. 2007) in a workflow managed by HTCondor software (Thain et al. 2005) quantified eleven ear and kernel traits from digital images as previously described (Miller et al. 2017). Six representative ears per plot were measured. Ear weight and grain weight was an average of the weight of the uppermost ear on the six representative plants in the plot and cob weight was measured on individual uppermost ears from the six representative plants in the plot (Table 1 and see Table S3). For all traits for which single plant measurements were taken, the same six representative plants were used for all measurements. See Table S2 for details on which traits were measured in each environment and Table S3 for raw phenotypic values.

**Table 1.** Summary of better parent heterosis (BPH) for 25 traits measured across 16 environments.

| Trait | Abbreviation | Locations | Plots | Average %BPH | % Entries with BPH<sup>a</sup> |
| --- | --- | --- | --- | --- | --- |
| Plant Height 14 DAP | PHt14 | 4 | 169 | 20.2 | 95.3 |
| Plant Height 21 DAP | PHt21 | 8 | 340 | 24.3 | 95.6 |
| Plant Height 28 DAP | PHt28 | 12 | 528 | 27.9 | 98.1 |
| Plant Height 35 DAP | PHt35 | 12 | 538 | 30.4 | 99.4 |
| Plant Height 42 DAP | PHt42 | 12 | 538 | 33.8 | 99.8 |
| Plant Height 49 DAP | PHt49 | 10 | 461 | 35.8 | 99.8 |
| Plant Height 56 DAP | PHt56 | 8 | 282 | 38.5 | 100.0 |
| Plant Height 63 DAP | PHt63 | 2 | 94 | 41.4 | 100.0 |
| Plant Height Maturity | PHt | 16 | 716 | 27.9 | 100.0 |
| Ear Height Maturity | EH | 16 | 716 | 36.5 | 99.3 |
| Days to Anthesis<sup>b</sup> | DTA | 16 | 670 | -4.4 | 82.1 |
| Days to Silk<sup>b</sup> | DTS | 16 | 670 | -5.0 | 87.0 |
| Leaf Number Above Ear | LNA | 12 | 538 | 7.6 | 53.9 |
| Leaf Number Below Ear | LNB | 8 | 375 | 3.1 | 20.0 |
| Cob Width | CW | 16 | 714 | 8.8 | 82.1 |
| Cob Length | CL | 16 | 714 | 20.3 | 93.1 |
| Cob Weight | CWT | 16 | 714 | 50.5 | 91.2 |
| Ear Width | EW | 16 | 710 | 13.5 | 95.4 |
| Ear length | EL | 16 | 713 | 18.6 | 91.6 |
| Ear Weight | EWT | 16 | 714 | 84.9 | 98.9 |
| Kernel Row Number | KRN | 16 | 713 | 7.2 | 58.6 |
| Kernel Height | KH | 16 | 710 | 26.1 | 89.6 |
| Kernel Width | KW | 16 | 709 | 8.7 | 59.4 |
| Kernel Depth | KD | 16 | 714 | 2.8 | 1.5 |
| Per Plant Grain Weight | GWT | 16 | 714 | 91.4 | 99.4 |

<sup>a</sup>Entry is the inbred/hybrid combination within an environment
<sup>b</sup>For these traits the better-parent performance was the lower value.

### Statistical Analyses

Better-parent heterosis (BPH) and percent better-parent heterosis (%BPH) were calculated for each trait and hybrid within each replicate block as BPH = hybrid phenotype - better-parent phenotype and %BPH = ((hybrid phenotype - better-parent phenotype)/Better-parent phenotype) x 100, respectively and then averaged across replicates within an environment. The average %BPH of the two replicates in each environment was used for subsequent analyses. For all traits except flowering time the higher parent was considered the better parent. For flowering time, the earlier parent was considered the better parent.

A mixed model analysis was performed using PROC GLM in SAS 9.0 (SAS Institute 2002) to partition variation into genotype, environment, genotype-by-environment interaction, and error variances for each trait with all sources of variation considered random. This analysis was done for inbred traits per se, hybrid traits per se, and heterosis for all 25 traits. Pearson correlation coefficients and corresponding significant tests was conducted using PROC CORR in SAS 9.0 (SAS Institute 2002). A mixed linear model was constructed by PROC MIXED in SAS 9.0 (SAS Institute 2002) to get the best linear unbiased prediction for each hybrid and inbred across the 16 environments: *y*_*i*_ = *μ* + *f*_*i*_ + *e*_*i*_ + *ε*_*i*_, where *y*_*i*_ is phenotypic value of individual *i*, *μ* is the phenotypic mean of multiple environments, *f*_*i*_ is genotype effect, *e*_*i*_ is environmental effect, and *ε*_*i*_ is the residual effect. All the variables except μ were considered as random effects (Bernardo 1994, 1996; Henderson 1975, 1984). The coefficient of variation (CV) for traits was calculated as the standard deviation divided by the plot mean. This is the most widely used parameter to quantify variability of traits with different units of measurement among individual plants and across environments (Munaro et al. 2011a).

### Statement on data and reagent availability

All raw phenotypic data is available in Supplemental Table 3 and GPS coordinates of locations and growth conditions are available in Supplemental Table 2.

## RESULTS AND DISCUSSION

### Better parent heterosis is variable across traits, environments, and developmental time

The majority of 25 measured traits exhibited significant genotype, environmental, and genotype-by-environment interaction effects in both the inbred lines and the hybrids across the 16 environments as well as for BPH (see Table S4). Better-parent heterosis (BPH) was detected for most of the 25 traits, and 16 of them exhibited BPH in more than 90% of hybrids (Table 1). Only two traits, leaf number below the ear (LNB) and kernel depth (KD), exhibited BPH in fewer than 50% of the hybrids. The average BPH varied substantially among the different traits (Table 1). Some traits such as grain yield per plant (GWT) exhibited BPH values greater than 90% BPH while other traits such as flowering time (DTA/DTS) exhibited a lower magnitude of BPH (-4.4 to -5.0%). However, for both of these traits the majority of hybrids exhibited BPH across all environments (Table 1).

The correlation of BPH across the traits studied varied substantially ranging from (*r*=-0.33 for EWT and PHt63 to 0.99 for EWT BPH and GWT BPH; Figure 1A and 1B), similar to previous observations (Flint-Garcia et al. 2009). A network visualization of the correlations between BPH for distinct traits revealed several trends (Figure 1B). Strong positive correlations were observed within groups of traits that likely share a common genetic, physiological and developmental basis, including yield related traits (cob, ear, and kernel traits) and vegetative traits including plant height at 14DAP through maturity and ear height (Figure 1A and 1B). Days to anthesis (DTS) BPH and plant height 63 days after planting (PHt63) BPH had strong negative correlations with grain weight (GWT) BPH, ear weight (EWT) BPH and ear width (EW) BPH because the better parent was the one that flowered early, and the hybrids flowered generally earlier than the better parent. PHt63 BPH was correlated with both vegetative and reproductive plant traits, connecting the two subgroups (Figure 1B).

**Figure 1.**
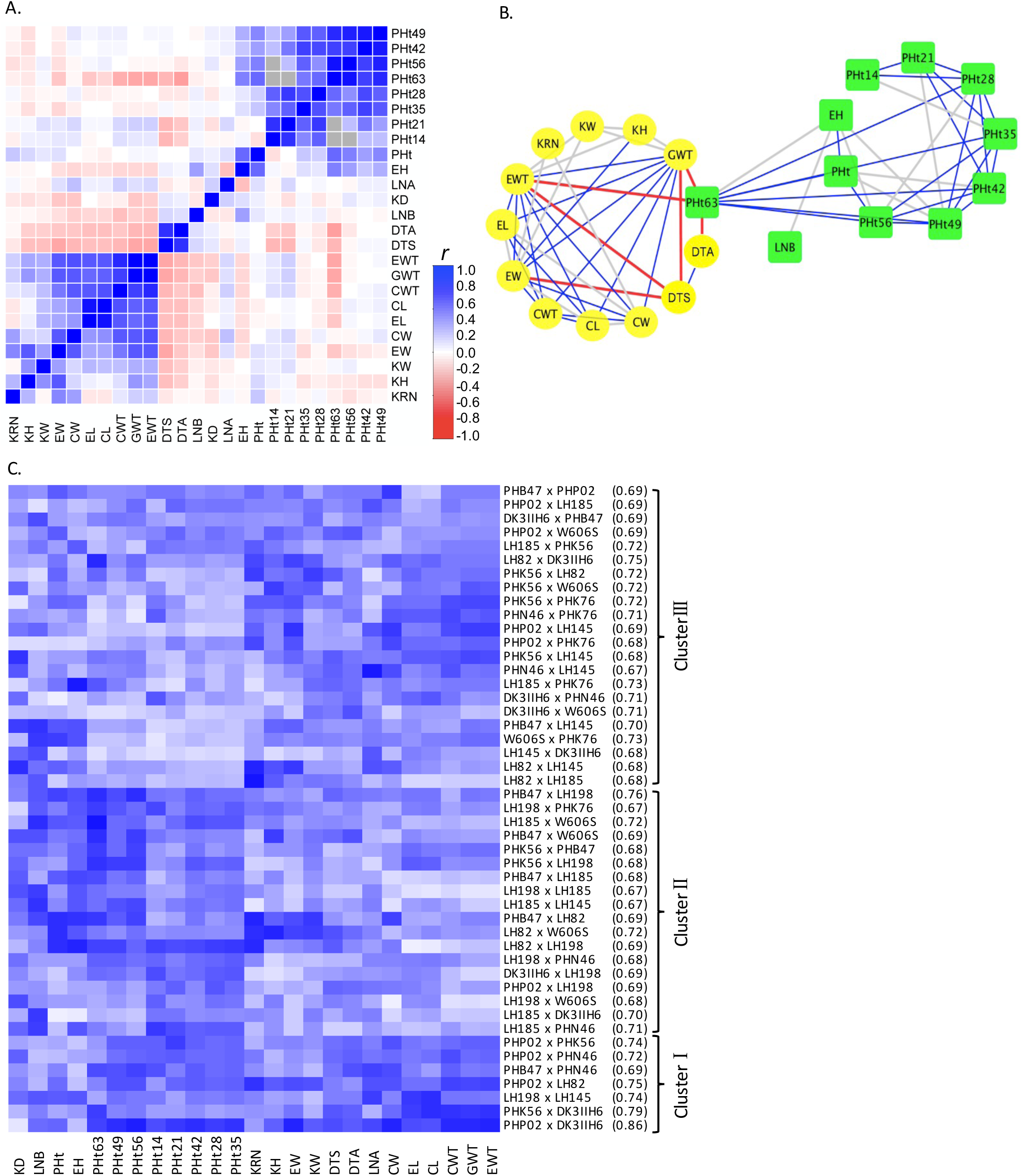
Better parent heterosis (BPH) comparisons for 25 traits and 47 hybrids across 16 environments. A) Pearson correlation coefficients (*r*) of BPH between traits; gray shaded cells indicate missing data. B) Network visualization of Pearson correlation coefficients of BPH between traits. Only correlation coefficients less than -0.3 or greater than 0.3 are shown. Traits in yellow circles and green rectangles are reproductive and vegetative traits, respectively. Red line, *r*<-0.3; gray line, 0.3<*r*<0.5; blue line, *r*>0.5. C) Average BPH rank scaled with white (highest BPH rank) to dark blue (lowest BPH rank). Hybrid genotypes are followed by the parental identity by state value.

The 47 hybrids could be assigned into three general clusters based on their relative heterosis performance (rank number) across the 25 traits (Figure 1C). The first cluster (n=7) exhibited consistently lower BPH for all the traits relative to other hybrids and was significantly enriched (Table S5) for within heterotic group hybrids (NSS x NSS). Hybrids in the second cluster (n=18) showed relatively high heterosis for yield-related and flowering time traits, but lower heterosis for most of the vegetative traits, while hybrids in the third cluster (n=22) were the opposite (Figure 1C).

Among the 47 hybrids genotypes, identity by state values ranged from 0.67 to 0.86 for the widest and narrowest crosses. Genotype clusters 1, 2, and 3 had identity by state averages of 0.755, 0.694, and 0.702 respectively. The hybrid genotypes in cluster 1, which had the lowest relative heterosis across all traits, was composed of relatively narrow crosses. The hybrid genotypes in cluster 2, which had the highest amount of heterosis across yield-related and flowering traits, was composed of relatively wide SSS x NSS crosses. This supports the historical convention of breeders crossing between heterotic pools of unrelated inbreds to maximize heterosis for yield related traits.

### There is low predictive capacity of heterosis over developmental time

It is desirable to identify traits early in development that predict heterosis and yield at the end of the season. Previous reports indicate that traits measured at maturity showed the highest level of heterosis (Falconer and Mackay 1996). However, it has been shown that heterosis could already be detected during early stages of maize seedling growth (Hoecker et al. 2006) and embryo development (Paschold et al. 2010; Meyer et al. 2007).

To determine the potential of heterosis based on early developmental stages to predict heterosis at later development stages we measured heterosis for plant height at seven developmental stages ranging from 14 days after planting to anthesis. A low correlation was observed between heterosis for plant height at early developmental stages and at anthesis (Figure 2). However, final plant height was more highly correlated with measures at developmental stages closer to anthesis. Overall, our results indicate that heterosis measured at the seedling stage is not predictive of heterosis at the adult stage.

**Figure 2.**
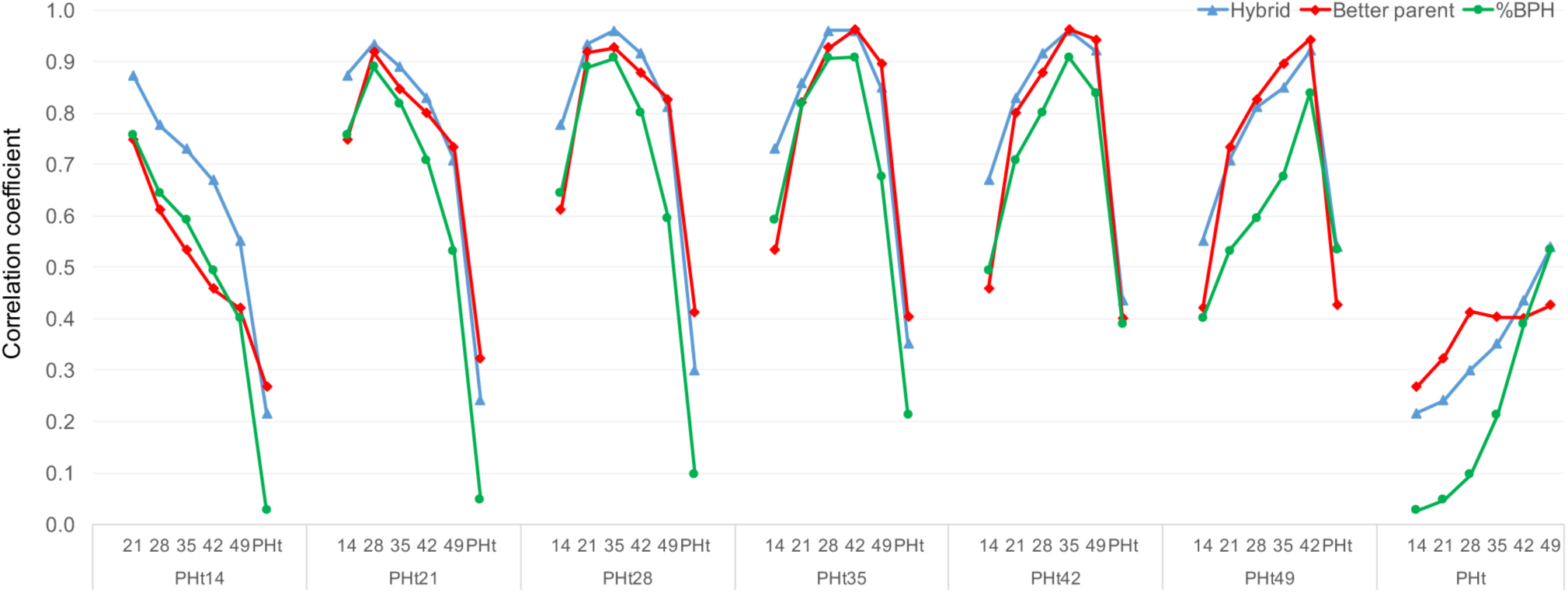
Correlation coefficient for percent better parent heterosis (%BPH), hybrid performance, and better-parent performance of plant height at different development stages in different environments. The numbers of 14-49 in x-axis indicate days after planting and PHt is plant height at physiological maturity.

These low levels of correlation could potentially be a product of low correlation for the hybrid performance, the better parent performance, or both. To evaluate what drives this reduced correlation in heterosis over increased windows of developmental time, correlation coefficients for hybrid performance over time and inbred performance over time were overlaid with heterosis correlations (Figure 2). Both hybrid performance and inbred performance showed a similar tendency over time, indicating that both hybrid and better parent performance have a comparable effect on the lack of correlation from early stages of development to maturity. Given the inability to predict heterosis levels, or even heterosis ranks, for the same trait (plant height) collected at different stages of development it is likely to be quite difficult to predict adult plant traits from seedling traits or to relate specific heterosis mechanisms observed in the seedling to those contributing to variation in heterosis for traits at maturity.

### Performance of hybrids is more stable than inbred lines within but not among environments

Differential responses of maize hybrids and/or inbred lines to environmental stimuli will result in altered levels of heterosis across environments (Munaro et al. 2011b; Munaro et al. 2011a). Evidence from multiple species indicates that hybrids performance is more stable across environments than inbred performance (Cole et al. 2009). This observation is consistent with the concept of “buffering” in which heterogeneous populations or heterozygous individuals are more stable than homogeneous populations or homozygous individuals (Allard and Bradshaw 1964; Schnell and Becker 1986; Cole et al. 2009). We compared the stability of inbred and hybrid traits both within an environment and among environments.

To evaluate stability across traits the coefficient of variation (CV) was used. The within plot CV for inbred lines in this study was greater than the within plot CV for hybrids for nearly every trait measured (Figure 3A), providing evidence for greater variability of inbred lines within environments for most traits. We also assessed the CV among environments for each trait in the inbred and hybrid lines (Figure 3B). For ten of the traits the inbred lines exhibited a significantly higher CV than the hybrids, indicating that for these traits instability across environments was driven more by the instability of inbred lines. However, for flowering time traits (DTA and DTS), hybrids had significantly higher CV than inbred lines across the environments. The remaining 13 traits did not exhibit significant differences between the hybrid and inbred lines for the CV among environments. For the plant height measurements over developmental time, the CV among environments decreased throughout time for both inbred lines and hybrids, indicating increasing stability of both hybrids and inbred lines at later developmental stages.

**Figure 3.**
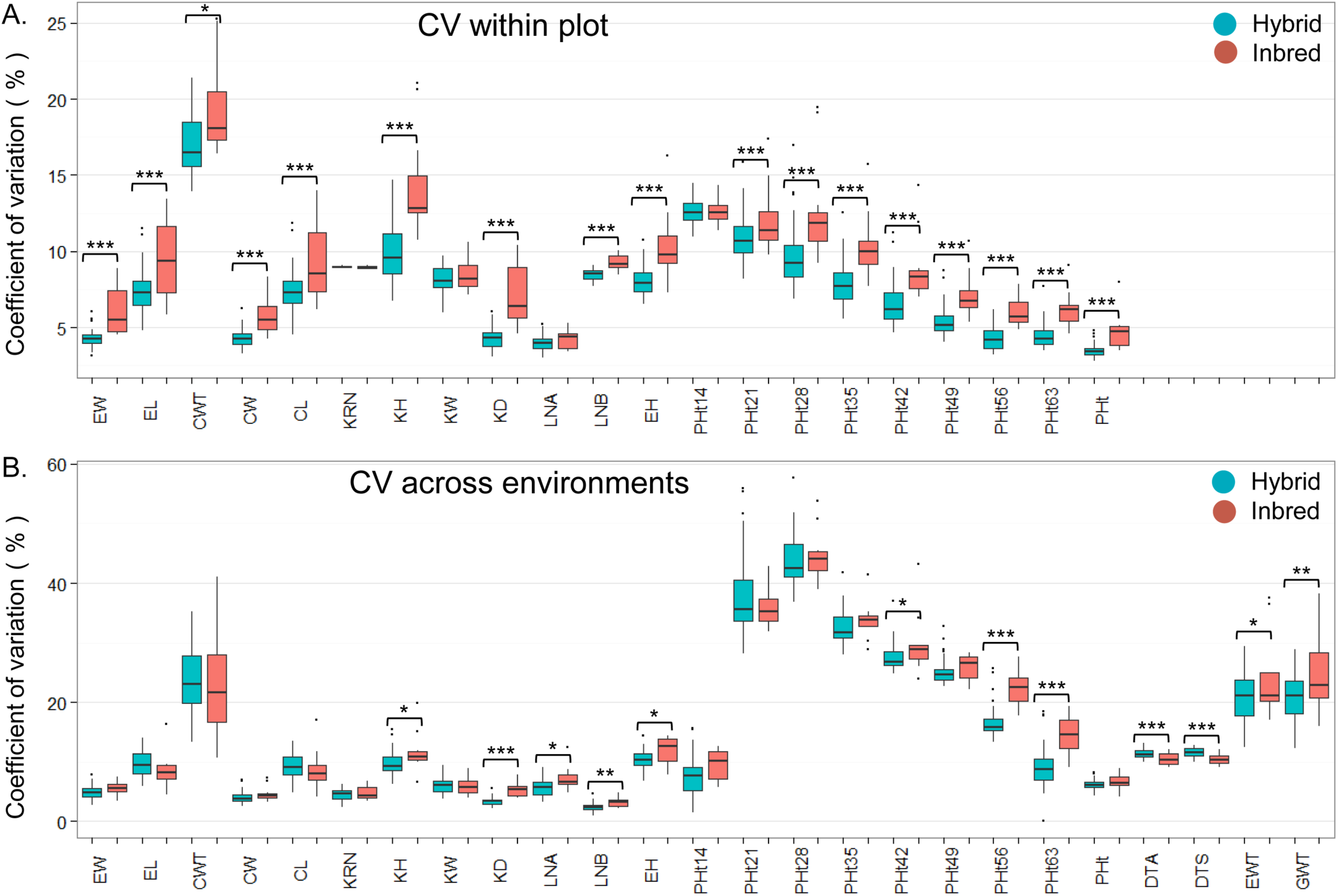
Coefficient of variation within and across environments for hybrid and inbred genotypes. A) Coefficient of variation within plot (6 plants were phenotyped within each plot). B) Coefficient of variation across all available environments for each trait. In each figure blue and red colors indicate hybrid and inbred, respectively. BLUP values of all available environments for each hybrid and inbred were used. * significant at *p*=0.05; ** significant at *p*=0.01; *** significant at *p*=0.001 in a two-tail t-test between the inbred and hybrid genotypes.

### Factors influencing environmental variation for heterosis are variable across traits

We were interested in assessing the factors contributing to the significant genotype-by-environment interaction effect on heterosis for most of the traits in this study. We focused on grain weight (GWT) and plant height at maturity (PHt). These traits have variable heritability, and BPH for these traits were not significantly correlated across genotypes or environments (Figure 1).

There were differences in the patterns of BPH among environments observed for GWT and PHt (Figure 4A and 4C). GWT generally expressed a greater BPH in low planting density environments, while planting density seemed to have little impact on BPH of PHt. For GWT, the correlation of IBS and BPH at high density was slightly higher (r=-0.58***) than for low density (r=-0.52***) indicating that BPH may be more affected by IBS at high density environments. However, both are highly correlated and IBS is affecting BPH under both conditions. For each of the traits the stability and average BPH was quite variable among hybrids (Figure 4). The hybrid that expressed the lowest and highest BPH based on BLUP values across all the environments were identified for each trait (indicated by arrows in Figure 4A and 4C). The stability of heterosis in these hybrids was evaluated across environments. Interestingly, for PHt the hybrid with the highest BPH exhibited consistently high levels of BPH while the hybrid with the lowest average BPH exhibited quite variable heterosis among environments (Figure 4D). However, this hybrid also had lower hybrid performance and therefore this result may be due to sensitivity to variable neighbor effects. The opposite pattern was observed for the hybrids with highest and lowest average BPH for GWT (Figure 4C). This trend was consistent across the entire set of 47 hybrids (see Figure S1). This may suggest that hybrids with the highest potential for GWT are the most responsive and have the potential to take advantage of favorable environments.

**Figure 4.**
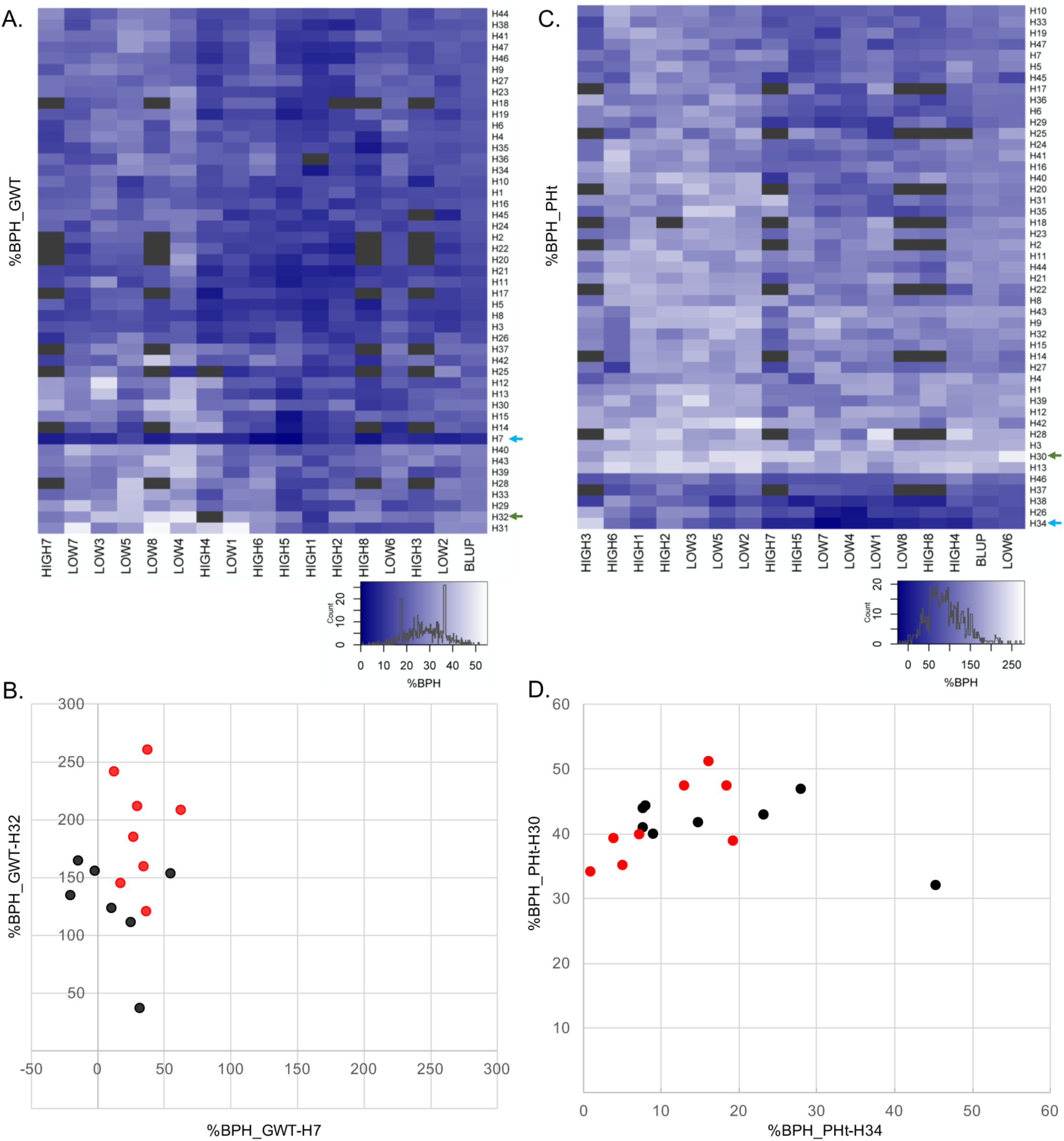
Percent better parent heterosis (%BPH) for grain weight (GWT) and plant height at maturity (PHt) for 47 hybrids across 16 environments. A and C) Heatmap of %BPH for GWT (A) and PHt (C); black shaded cells indicate missing data. The green and blue arrow in each plot indicates the hybrids that have the highest and lowest %BPH across all 16 environments based on BLUP values. Environments and hybrids were clustered using hierarchical clustering (trees not shown). B and D) Highest (indicated by green arrows in A and C) vs. lowest (indicated by blue arrows in A and C) %BPH hybrids across all environments for GWT (B) and PHt (D). Red dots are the eight low-density environments and black dots are the eight high-density environments. H7 is PHP02 x DK3IIH6, H30 is LH185 x DK3IIH6, H32 is LH198 x LH185, H34 is LH82 x W606S.

BPH is a measure of the difference in performance of the hybrid relative to the parents. Environmental influences on BPH could reflect changes in hybrid performance, changes in inbred performance or a combination of both. We investigated the patterns of BPH, hybrid performance and inbred performance for GWT and PHt in a selected set of hybrids (Figure 5). We first assessed the patterns for the hybrids with the highest average BPH for GWT (Figure 5A) and PHt (Figure 5E). We also assessed the patterns for the hybrid with the greatest (Figure 5B and 5F) or least (Figure 5C and 5G) standard deviation for BPH ranks among the environments. These reveal a variety of patterns in the trend of inbred and hybrid performance relative to variation in BPH values among environments. There are examples, such as Figure 5E, in which the reduction in heterosis in some environments is due to reduced hybrid performance with relatively stable inbred performance. In other examples, such as Figure 5C, the changes in heterosis seem to be driven by changes in the inbred performance among the environments.

**Figure 5.**
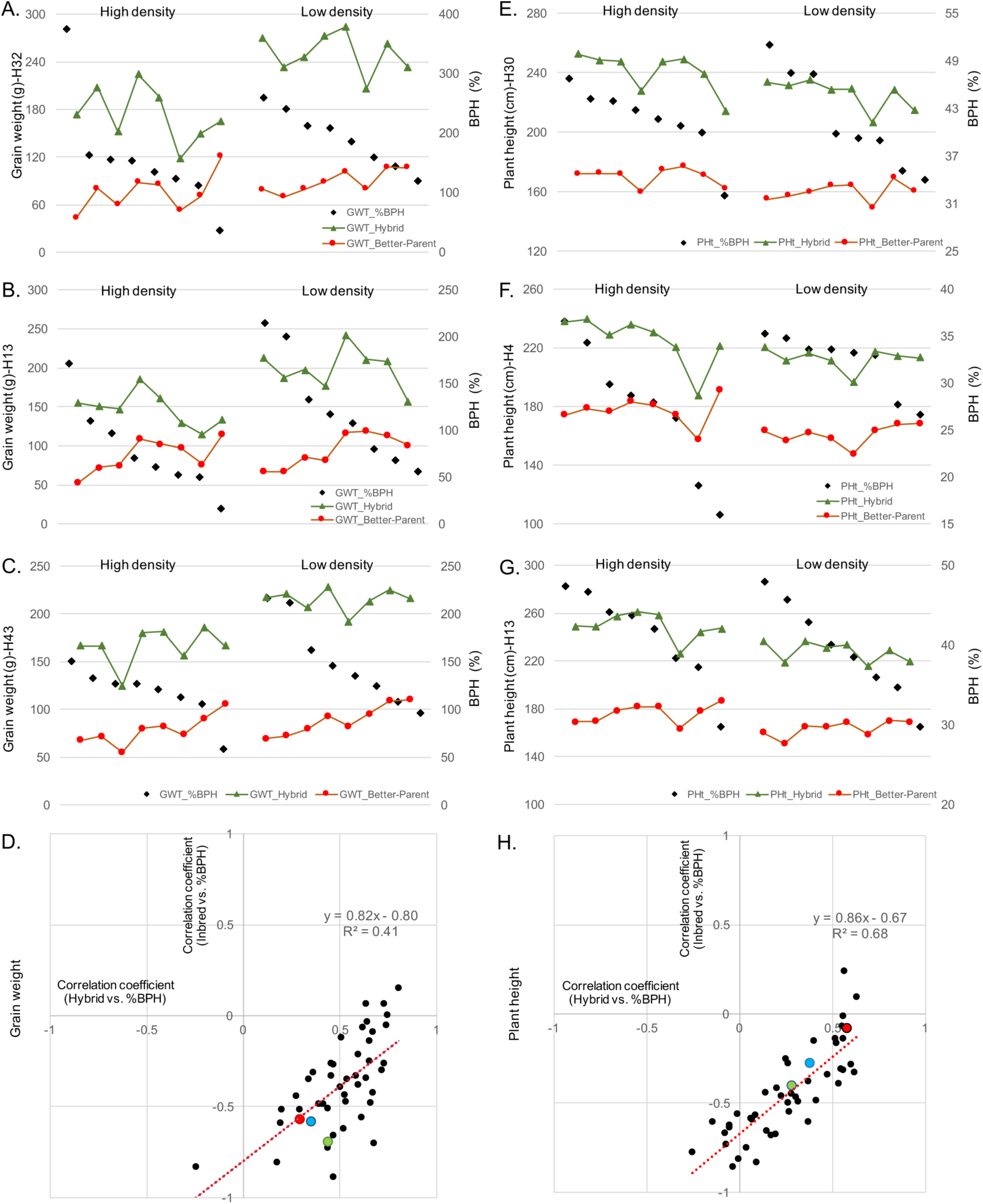
Relationships among percent better parent heterosis (%BPH), hybrid, and better-parent performance. Plots A-D are for grain weight (GWT) and E-H are for plant height at maturity (PHt). A and E) Hybrids with the highest %BPH across 16 environments. B and F) Hybrids with the highest standard deviation of the rank of %BPH among all 47 entries. C and G) Hybrids with the lowest standard deviation of the rank of %BPH among all 47 entries. D and H) Correlation coefficient of hybrid vs. %BPH and better-parent vs. %BPH (BLUP value across 16 environments for each hybrid). Colored dots represent the highest %BPH (red – A and E), highest standard deviation of the rank of %BPH (green – B and F), and lowest standard deviation of the rank of %BPH (blue – C and G). For A-C and E-G dots along the x-axis represent each of the 16 environments.

We proceeded to assess the relative contribution of variation in the inbreds and hybrids to the environmental variation for BPH for all 47 hybrids for GWT and PHt (Figure 5D and 5H). The correlation of better parent performance and BPH (y-axis) was plotted against the correlation of hybrid and BPH (x-axis). As expected, the performance of the better parent tends to be negatively correlated with heterosis while the performance of the hybrid is positively correlated with heterosis. If variation for better parent performance and hybrid performance equally contribute to heterosis variation we would expect a slope of one in the regression line of this plot. The observed slope was less than one, indicating that variation in the hybrids was contributing slightly more to the observed BPH values than variation in the inbred lines. There are differences in the distribution of the correlation values for GWT (Figure 5D) and PHt (Figure 5H). For GWT, 46 of 47 hybrids have a positive correlation between hybrid performance and heterosis (Figure 5D) suggesting that heterosis for GWT is influenced by hybrid performance in all genotypes. In contrast, there are a number of hybrids without significant correlations between hybrid performance and heterosis for PHt (Figure 5H).

We assessed the relative influence of better parent and hybrid variation on BPH for all 25 traits measured in this study (Table S6). In the majority of cases the hybrid performance is positively correlated with heterosis while the better parent performance is negatively correlated with heterosis. However, the relative strength of the correlations varied among different traits. For traits such as KD, PHt, DW there was a much stronger correlation between better parent performance and heterosis. Environmental variation for heterosis for other traits such as CWT, KW, and EL are more strongly influenced by the hybrid performance (Table S6). Interestingly, GWT showed equal strength of correlation for both hybrid performance with heterosis and better parent performance with heterosis. There was, however, a significant negative correlation between “noise” (residual from ANOVA using BPH) and the correlation of better parent performance and BPH (r=-0.77***), which may impact the ability to accurately assess the relative contribution of inbreds and hybrids to observed BPH.

Corn yields have increased continuously since hybrids were first commercially grown in the 1930s. However, the increase in yield of commercial hybrids has not been attributed to an increase in heterosis (Fasoula and Fasoula 2005). Indeed, the percentage of heterosis has not changed substantially, and by some estimates has decreased slightly over time due to the higher percentage rate of gain in yield for inbred lines relative to hybrids (Duvick 1999; Troyer and Wellin 2009). Our data suggest that variation in the performance of inbred lines and hybrid lines in different environments will influence the amount of heterosis. The relative influence of hybrid variation and inbred variation on heterosis is variable across the traits that were measured in this study. It is worth noting that in some extreme environments inbred lines may be severely affected while hybrids are not, and this outcome will influence measures heterosis (Griffing and Zsiros 1971). However, in the moderate environments surveyed in this study we find important contributions of both hybrid and inbred performance to heterosis variation.

## ACKNOWLEDGEMENTS

We thank Peter Hermanson, Brad Keiter and James Satterlee for technical assistance. This work was supported in part by the Minnesota Corn Research and Promotion Council (Project Number 4108-16SP), the Minnesota Agricultural Experiment Station (Project 13-014), the Iowa Corn Growers, and Iowa State University’s Plant Sciences Institute.

